# Novel Organ-Specific Genetic Factors for Quantitative Resistance to Late Blight in Potato

**DOI:** 10.1101/567289

**Authors:** Deissy Katherine Juyo Rojas, Johana Carolina Soto Sedano, Agim Ballvora, Jens Léon, Teresa Mosquera Vásquez

## Abstract

Potato, *Solanum tuberosum*, is one of the highest consumed food in the world, being the basis of the diet of millions of people. The main limiting and destructive disease of potato is late blight, caused by *Phytophtora infestans*. Here, we present a multi-environmental analysis of the response to *P. infestans* using an association panel of 150 accessions of *S. tuberosum* Group Phureja, evaluated in two localities in Colombia. Disease resistance data were merged with a genotyping matrix of 83,862 SNPs obtained by 2b-restriction site–associated DNA and Genotyping by sequencing approaches into a Genome-wide association study. We are reporting 16 organ-specific QTL conferring resistance to late blight. These QTL explain from 13.7% to 50.9% of the phenotypic variance. Six and ten QTL were detected for resistance response in leaves and stem, respectively. *In silico* analysis revealed 15 candidate genes for resistance to late blight. Four of them have no functional genome annotation, while eleven candidate genes code for diverse proteins, including one leucine-rich repeat kinase.

## 1. Introduction

Potato (*Solanum tuberosum*) is the most important non-cereal crop and plays a basic role for the food security and nutrition [1]. Also, potatoes are an important crop for farmers and consumers because of its ability to efficiently transform inputs into a high-quality food with an excellence consumer acceptance.

The predominately cultivated potato species worldwide is *S. tuberosum* L. (2n=4x=48), this species contains two groups: Chilotanum (tetraploid potato) and Andigenum [2]. The Group Andigenum has a subset of diploid potatoes (2n=2x=24), the Group Phureja [3]. Phureja has been used worldwide as a genetic source and has contributed to understand the genetic basis of several agronomic traits of potato. It is worth to notice that the current potato reference genome sequence is obtained from DM1-3516 R44 a Phureja genotype [4].

Potato, as any other vegetative multiplied crop, is affected by several pests and diseases and the most devastating disease affecting this crop is late blight caused by the oomycete *Phytophthora infestans* [5–8]. Late blight has been reported in the main regions where potato is grown, including more than 130 countries distributed over Asia, Africa, Oceania, Europe, South, Central and North America [9]. Thereby, great concern exists worldwide for possible emergence of new *P. infestans* strains as well as for the number of countries affected by the disease that could be increasing. Although chemical control of late blight is possible, the best alternative is through the use of resistant varieties. In potato, both, pathotype-specific (vertical resistance) and race non-specific resistance (field, horizontal, polygenic or partial resistance) have been reported [6,7,10–12].

The potato breeding programs worldwide are focused to obtain late blight resistant cultivars though conventional breeding [13–15]. Despite these programs have successfully developed partial resistant cultivars [16], several challenges remain to be faced. If the selection is based on vertical resistance exists the risk of its breakdown caused by an eventual emergence of new *P. infestans* races [11], but if the selection is based on horizontal resistance, this is highly dependent on the environmental conditions, as well as on the pathogen strains. Thus, the cultivar resistance could change depending on the geographic region [17]. To face these challenges, it is important the detection of quantitative trait loci (QTL) that govern late blight resistance in both tetraploid and diploid potatoes, through methods based on genetic variants capture by DNA molecular markers such as Genome-wide association studies (GWAS) and on multi-environmental phenotyping. This will reduce the time required in the process, and also will contribute to a better understanding of the genetic basis of potato - *P. infestans* interaction.

In potato, several successful association mapping studies have been achieved for traits such as diseases resistance [12,18–20]. For diploid potatoes, the search for sources of resistance to late blight through GWAS has been relatively scarce. Two candidate genes and three SNPs associated with quantitative resistance to late blight in the Group Phureja were identified using 371 SNP distributed along the genome [18,21].

In the polygenic resistance one of the major limitations is the characterization of the plant-pathogen interaction due to many influencing factors, including the biology, genetics and evolution of the pathogen in addition to the environmental conditions [22]. To facilitate the understanding of polygenic resistance, it has been proposed to subdivide the behavior of the disease in the different affected organs of the plant [23].

Studies indicate that differences in organ-specific resistances are resulting in an increase of transcription levels, although there is also the possibility that organs develop hypersensibility response (HR) [23–25].

The aim of the presented study is to identify QTL for resistance to late blight in landrace accessions *S. tuberosum* Group Phureja. We employed a GWAS using phenotypic data from the characterization of disease resistance conducted in two localities and genotyping data obtained from GBS and 2b-RAD approaches. Here, we report 16 organ-specific QTL for resistance to late blight in *S. tuberosum* Group Phureja. The analysis of DNA sequence corresponding to the QTL reveals 15 candidate defense genes that both, contribute to a better understanding of the genetic control of late blight resistance in diploid potato, and also could be used in the future in potato breeding programs.

## 2. Materials and Methods

### 2.1. Plant material

The association panel consists of 150 accessions of *S. tuberosum* Group Phureja from the Working Collection of the Potato Breeding Program at Universidad Nacional de Colombia. From them, 109 are landrace accessions belonging to the Colombian Core Collection (CCC). Five are Colombian commercial genotypes, denoted as commercial cultivars, Criolla Colombia (CrCol), Criolla Guaneña (CrGN), Criolla Latina (CrLSE1), Criolla Paisa (CrPSE1), and Criolla Galeras (CrGL). Four are diploid accessions from the *IPK Germplasm Bank of Germany* (Leibniz-Institut für Pflanzengenetik und Kulturpflanzenforschung), which correspond to materials collected in Colombia, Peru and Bolivia by the International Potato Center (CIP). Three accessions are from a segregating F1 population derived from a cross between the susceptible parental 48A.3 and the resistant parental 2A4, that exhibit different levels of resistance to *P. infestans*, and 20 are a new set of landrace accessions (CA) that were collected in Nariño province, located at south of Colombia, in 2013. The cultivars ICA Unica and Diacol Capiro were the resistant and the susceptible controls to late blight, respectively.

Each accession was multiplied vegetatively, late blight-free, using seed potato tubers at the experimental station, Centro Agropecuario Marengo of Universidad Nacional de Colombia, with an altitude of 2,516 meters above sea level (masl), annual average temperature of 13.7 °C, and cumulated rainfall of 669.9 mm/year. The production of seeds was carried out during three consecutive crop cycles under the Colombia traditional conditions of potato crop [26].

### 2.2. Genotyping approaches

GBS libraries were prepared and analyzed at the Génome Québec Innovation Centre from McGill University, while the 2b-RAD was conducted at the Agriculture and Agri-Food Canada (AAFC). The *Pst*I*/Msp*I enzymes were used for GBS digestion and the selective amplification was performed according to [27]. The *Alf*I enzyme was used for 2b-RAD approach. The DNA fragments generated were sequenced using the MiSeq system from Illumina®.

DNA sequencing raw data from GBS and 2b-RAD were analyzed for SNP polymorphisms detection, according to the NGSTools Eclipse Plugin (NGSEP) [28]. The SNP calling was performed by alignments to the current potato reference genome from *S. tuberosum* Group Phureja clone DM1-3 (v4.03) [29], in order to obtain an annotated SNP matrix.

The filters applied to the annotated SNP matrix allowed to select those SNPs with quality score higher than 40 (-q 40), 0.05 Minor Allele Frequency (MAF), minimum distance of 5 bp between variants, and SNPs located in non-repetitive regions according to the RepeatMasker annotated repeats [29]. Then, the SNP matrix was filtered using SAS® software (SAS institute Inc.). The criteria such as a MAF higher than 0.05 and less than 10% of distorted molecular markers were allowed and those SNPs with Mendelian segregation of 1:1 were selected.

### 2.3. Evaluation of potato response to late blight disease

The evaluation of the response of the association panel to late blight was carried out in two localities during different crop cycles, starting in May 2010 until July 2014. The combination of localities and the crop cycles conformed seven environments, with specific conditions, that were coded ENV1 to ENV7 (Table 1). The localities were, La Unión in Antioquia, (5°58′22″N 75°21′40″O) and Subachoque, in Cundinamarca (4° 55′ 41″ N, 74° 10′ 25″ W). These localities were selected as important potato production regions in Colombia, and have a high natural inoculum pressure of *P. infestans*. The experimental design used for the phenotypic evaluation consisted of an incomplete block design with three biological replicates (plant clones) per genotype, distributed in blocks of 12 experimental units of three tubers. The controls for resistance and susceptibility were the cultivars Ica Unica and Diacol Capiro, respectively [30]. The plants were evaluated for their response to *P. infestans* under natural infection conditions and inoculum pressure of RG57 (EC-1 clonal lineage), mating type A1, which is the predominant pathogen presented in both localities [31].

**Table 1.**
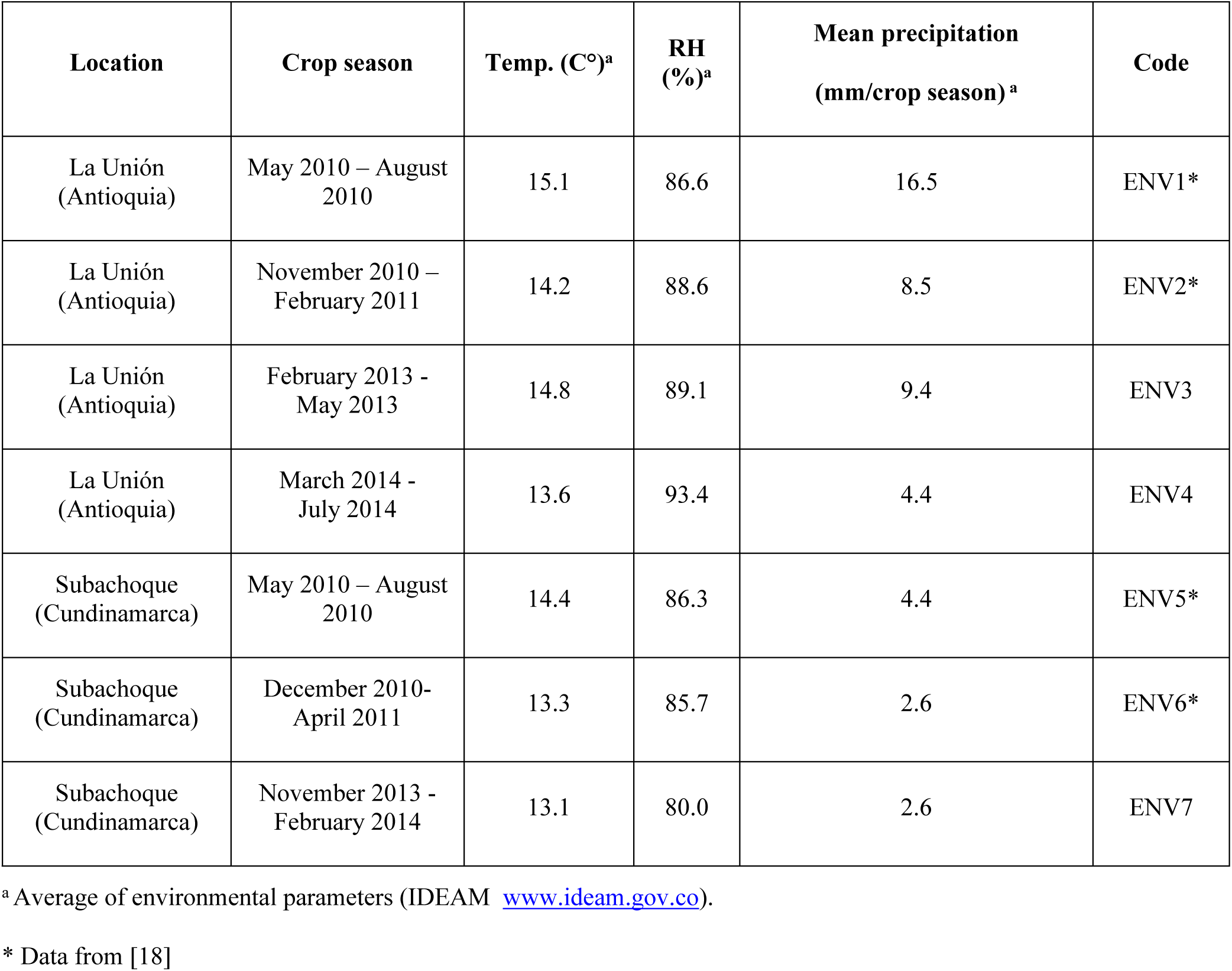
Description of each environment for the phenotyping evaluation for resistance to Phytophthora infestans in Group Phureja. It is shown the name of location, the crop season, mean temperatures, mean relative humidity (RH), mean precipitation and code.

| Location | Crop season | Temp. (C°)<sup>a</sup> | RH (%)<sup>a</sup> | Mean precipitation (mm/crop season)<sup>a</sup> | Code |
| --- | --- | --- | --- | --- | --- |
| La Unión (Antioquia) | May 2010 – August 2010 | 15.1 | 86.6 | 16.5 | ENV1* |
| La Unión (Antioquia) | November 2010 – February 2011 | 14.2 | 88.6 | 8.5 | ENV2* |
| La Unión (Antioquia) | February 2013 - May 2013 | 14.8 | 89.1 | 9.4 | ENV3 |
| La Unión (Antioquia) | March 2014 - July 2014 | 13.6 | 93.4 | 4.4 | ENV4 |
| Subachoque (Cundinamarca) | May 2010 – August 2010 | 14.4 | 86.3 | 4.4 | ENV5* |
| Subachoque (Cundinamarca) | December 2010- April 2011 | 13.3 | 85.7 | 2.6 | ENV6* |
| Subachoque (Cundinamarca) | November 2013 - February 2014 | 13.1 | 80.0 | 2.6 | ENV7 |

<sup>a</sup> Average of environmental parameters (IDEAM www.ideam.gov.co).
\* Data from [18]

The severity of late blight was estimated and scored according to the Percentage of Direct Visual Estimation of the disease (PDVE) [32] in leaves and stems following the concepts presented by Bock et al., [34] and the procedure described by Álvarez et al., [18]. The evaluations were conducted by expert raters in late blight, who made the estimations along the experiments. The diseased leaves and stems were assessed independently per each plant and for each organ. The severity was estimated as the percentage of the affected area for leaves and stems through a visual direct estimation. For stems, the PDVE was averaged over the number of stems of each genotype. The evaluation started 40 days post sowing (dps) and were performed weekly until the susceptible control was completely affected (PDVE = 100%). Disease progress in time for each plant was analyzed by estimating the Area Under Disease Progress Curve (AUDPC) and the Relative Area Under Disease Progress Curve (rAUDPC) to the total area [33,35]. The rAUDPC value for each genotype per environment was calculated by using the PROCMEAN procedure of SAS®.

In order to evaluate the resistance response of the Phureja genotypes to late blight infection, previous data of the resistance in leaves in the population were taken into account. These data correspond to the phenotypic evaluation of late blight response of 104 diploid accessions of the Group Phureja (CCC), in the same two localities from May 2010 to February 2011 (ENV1, ENV2, ENV5, and ENV6) [18] (Table 1) and these evaluations were integrated for the analysis with new evaluations for the same plant material conducted from February 2013 until July 2014 (ENV3, ENV4, and ENV7).

### 2.4. Analysis of genetic stability

The GGE-biplot criteria were centered by two (centered G + GE), without scaling and singular value partitioning (SVP). GGE-biplot analysis was performed using the R package GEA-R [36].

### 2.5. Statistical analysis

The rAUDPC values were tested for variance components of genotype, genotype by environment, and experimental error through an ANOVA (SI 1). All analyses were performed using the SAS® software. AUDPC data for the trials of late blight resistance evaluation of the CCC population were included for best linear unbiased predictors (BLUPs) estimation and association analysis. These data correspond to the phenotypic evaluation of late blight disease response of 104 diploid accessions of the Group Phureja CCC, at La Unión from May to August 2010, from May to August 2010 at Subachoque, from November 2010 to February 2011 at La Unión, and from December 2010 to April 2011 at Subachoque [18] (Table 1).

### 2.6. Genome-wide association study for late blight resistance

GWAS was performed for marker main and marker by locality interaction effects simultaneously using a liner mixed model in PROC MIXED procedure in SAS® program (SAS version 9.2) following the model:

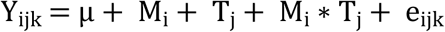

Where Y_ijk_ is the phenotypic value; µ is the general mean; M_i_ is the random effect of i-th marker genotype; T_j_ is the locality effect fixed of j-th locality; M_i_ * T_j_ is the interaction random effect of i-th marker with j-th local; and e_ijk_ is the residual. To determine traits of interest in the genome wide detection analysis, a log-of-odds (LOD) threshold with P-value ≤ 0.00001, and 1000 permutations was determined. The QTL model comprises an iterative multiloci procedure. Therefore, the most informative SNP (QTL) was set as a fixed factor during each calculation iteration step. All remaining markers were again incorporated in the next iteration round and re-analyzed. The starting point of each calculation round was determined by the result of the previous iteration. P-values of significant markers were corrected using Probability of False Discovery Rate (PFDR) of lower than 0.05, implemented in the SAS procedure PROC MULTTEST according to Benjamini and Yekutieli [37]. This procedure was repeated until no marker could be detected, which led to a reduction in significant markers and thereby a reduced number of false-positive QTL. A confidence interval of 1000 bp was chosen on both sides of the most significant SNP and designated as putative QTL. SNPs were combined to one joint QTL depending on their estimated (significant) P-value from the first iteration of the multi loci procedure. Therefore, the size of the genetic interval was dependent on the significance value of flanking SNPs.

The phylogenetic relationships of the 150 potato genotypes were determined based on the allele frequency of each SNP, the genotypes were assigned to an eventual population, using a Bayesian modeling in the software STRUCTURE v.2.3 [38]. The population structure matrix was used as the matrix Q in the association model.

### 2.7. QTL mapping and candidate gene proposed

The DNA sequences harboring SNPs identified as statistically associated with the phenotypic data of the response to late blight were explored through BLAST (Basic Local Alignment Search Tool) analysis to the current potato reference genome (v4.04). These sequences were searched also in the potato consensus QTL map [39] in order to find co-localizations of SNPs with previous reported QTL.

## 3. Results

### 3.1. GBS and 2b-RAD genotyping in *Solanum tuberosum* Group Phureja

Plant genome-wide genotyping for single nucleotide polymorphism (SNP) detection was performed by GBS [40] and 2b-RAD [41] approaches, using Illumina Hi-seq (Illumina, Inc.), which produced both in total 349,965,885 reads. Using GBS were obtained 302,965,885 reads of 90 bp, while with 2b- RAD 46,502,388 reads of 36 bp. On average, 88% of the reads, were aligned to unique positions in the potato reference genome (v4.04) [29]. Based NGSEP analyses a total of 87,656 SNPs were identified. From them, 85,755 SNPs were obtained from GBS and 1,901 from 2b-RAD. Finally, the subsequent filtering criteria reduced the matrix to 83,862 SNPs. These SNPs are distributed across of the 12 potato chromosomes with an average of 6,988 for each chromosome. The higher densities of SNPs were found in chromosomes I, III, and IV with 10,427; 8,237 and 8,160 SNPs, respectively (Figure 1). From the resulting matrix of 83,862 SNPs, 60% (49,948) corresponds to transitions while 40% (33,914) are transversions, with a ratio of 1.47 (SI 2).

**Figure 1.**
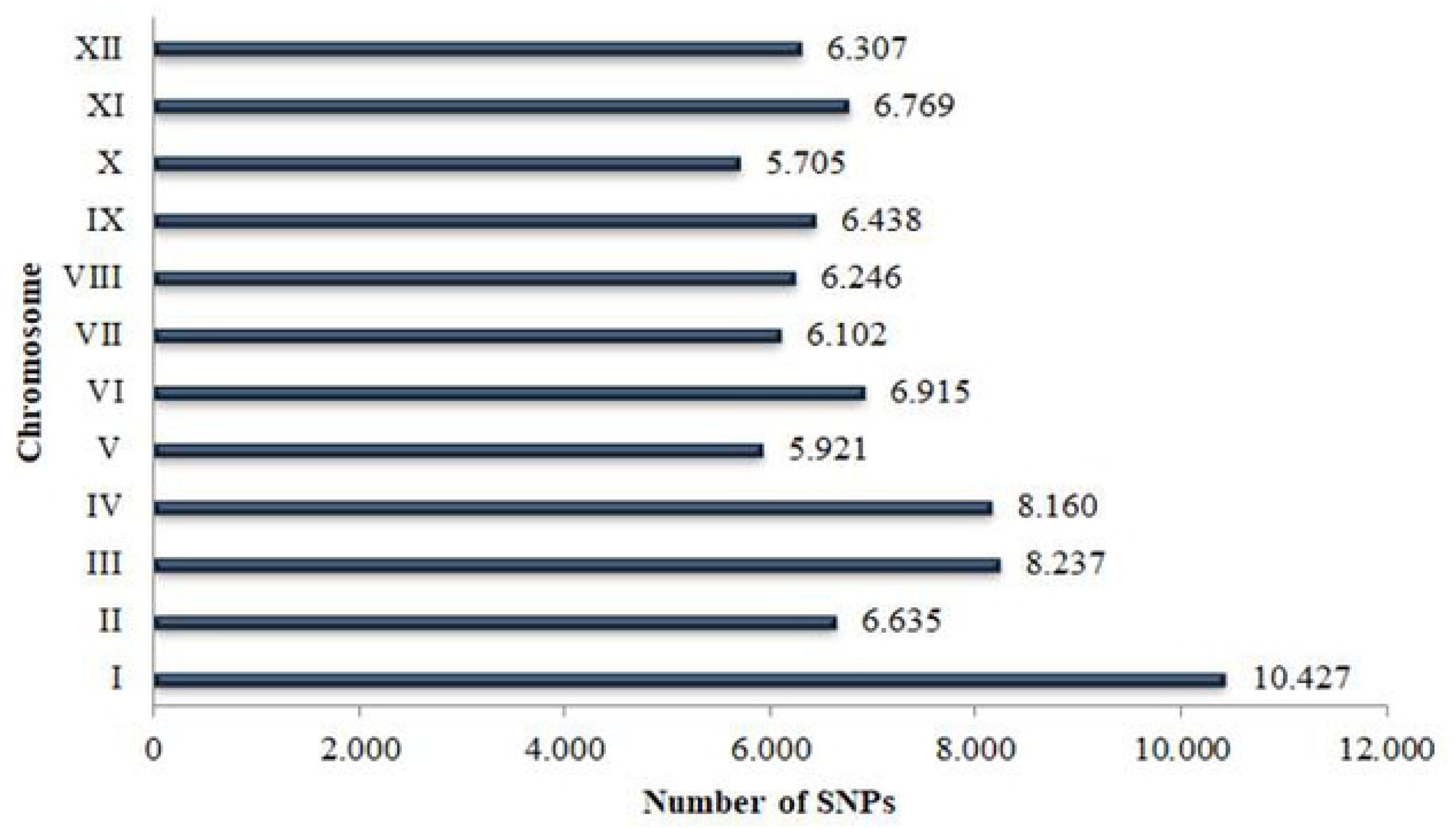
Number of SNPs mapped for each chromosome in the potato genome. Numbers on the right of bars show the total SNPs identified through GBS and 2b-RAD located in each chromosome as indicated by roman letters on y-axis.

### 3.2. Evaluation of late blight response in an association panel of Group Phureja

Both AUDPC and rAUDPC values for the genotypes for each locality and season showed a continuous distribution. The AUDPC values observed among the genotypes were the highest in the ENV6 (ranged from 1,400 to 13,325), followed by ENV5 (from 503.5 to 11,964.2) and then, by ENV2 (from 1,416.6 to 10,227.7). In the Table 2, it is shown the number of evaluated genotypes, the AUDPC and rAUDPC ranges.

**Table 2.** Distribution of the AUDPC and rAUDPC values in the Group Phureja association panel. Number of genotypes evaluated, range of AUDPC and rAUDPC values.

| Location/code | Evaluated genotypes | AUDPC range | rAUDPC range |
| --- | --- | --- | --- |
| ENV1 | 112 | 437.5 – 12,835 | 0 – 3.67 |
| ENV2 | 112 | 1416.6 – 10227.7 | 0.28 – 2.08 |
| ENV3 | 111 | 0 – 1456.6 | 0 – 0.46 |
| ENV4 | 114 | 0 – 834.1 | 0 – 0.23 |
| ENV5 | 112 | 503.5 – 11964.2 | 0.07 – 1.89 |
| ENV6 | 112 | 1,400 – 13,325 | 0.25 – 2.37 |
| ENV7 | 137 | 4.16 – 1713.4 | 0 – 0.40 |

### 3.3. Stability analysis

With the purpose of evaluate the performance of Phureja genotypes under different environments; a genotype x environment analysis was conducted through a GGE-biplot analysis of Multi-Environment Trials (MET). The rAUDPC values of each genotype in each plant organ (leaves and stems) were analyzed independently. The GGE-Biplot analysis shows that the two principal components explained 77.5% and 91.1% of the total variance caused by G_plant_ + G_pathogen_ + E for Phureja resistance to late blight in leaves and stem, respectively (Table 3 and Figure 2). The leaves analysis showed the two mega-environments conformation, MET 1 (ЕNV1, ЕNV2, ENV5, and ЕNV6) and MET 2 (ENV3, ENV4 and ENV7). But for stem analysis no mega-environments were detected. The rAUDPC values of the genotypes were able to discriminate among seven environments tested. The behavior of genotypes for both leaves and stem fall into two sectors of the graphic. For leaves resistance, the genotypes behavior seems to be quite similar in MET 1, where, the genotypes exhibit more susceptibility in the environment ENV1. In MET 2, the environments ENV3 and ENV4 show also the most susceptible genotypes (Figure 2).

**Table 3.** Summary of genome associations for resistance to late blight by location. Plant organ, localization, genome position, QTL for late blight resistance, its significance and gene annotation were the QTL resides.

| Plant organ | Location | Genome position | QTL name | P-value | R<sup>2</sup> | Gen annotation (Arabidopsis and Tomato) |
| --- | --- | --- | --- | --- | --- | --- |
| Leaves | Subachoque (Cundinamarca) | Chr 1 Pos 37547438 | RPiLSub1 | 6,2616E-08 | 18.8% | Protein Glycoside hydrolase, catalytic domain |
|  |  | Chr 3 Pos 54281967 | RPiLSub3 | 9,0818E-09 | 25.1% | Chloroplast protein with Zinc finger domain |
|  |  | Chr 6 Pos 40366150 | RPiLSub6 | 2,5174E-06 | 16.3% | Integral membrane transporter family protein |
|  | La Unión (Antioquia) | Chr 3 Pos 3162604 | RPiLLU3a | 9,15844E-11 | 28.5% | Ubiquitin receptor protein associated with PEX2 and PEX12. |
|  |  | Chr 3 Pos 54281967 | RPiLLU3b | 4,49269E-09 | 24.3% | Chloroplast protein with Zinc finger domain |
|  |  | Chr 11 Pos 7171775 | RPiLLU11 | 8,99280E-06 | 13.7% | Receptor-like protein kinase |
| Stem | Subachoque (Cundinamarca) | Chr4 Pos 9090691 | RPiSSub4 | 2,04E-06 | 17.3% | Leucine-rich repeat protein kinase family protein |
|  |  | Chr 6 Pos 11035921 | RPiSSub6 | 1,14E-09 | 25.8% | SIT4 phosphatase-associated family protein |
|  |  | Chr 7 Pos 53724116 | RPiSSub7 | 1,07E-09 | 25.8% | Actin filament-based movement. Endocytosis |
|  |  | Chr 10 Pos 51118650 | RPiSSub10 | 1,02E-07 | 20.8% | DHHC-type zinc finger family protein |
|  |  | Chr 12 Pos 57373550 | RPiSSub12 | 1,27E-08 | 23.2% | No functional annotation |
|  | La Unión (Antioquia) | Chr 5 Pos 45069210 | RPiSLU5 | 3,00E-08 | 22.2% | Ser/Thr protein kinase, BLUS1 |
|  |  | Chr 10 Pos 4646061 | RPiSLU10 | 3,53E-11 | 27.7% | No functional annotation |
|  |  | Chr 12 Pos 3076623 | RPiSLU12a | 1,60E-14 | 33.3% | Serine/threonine-protein kinase. Plasma membrane |
|  |  | Chr 12 Pos 38707075 | RPiSLU12b | 7,64E-23 | 50.9% | No functional annotation |
|  |  | Chr 12 Pos 56839213 | RPiSLU12c | 6,32E-18 | 38.4% | No functional annotation |

**Figure 2.**
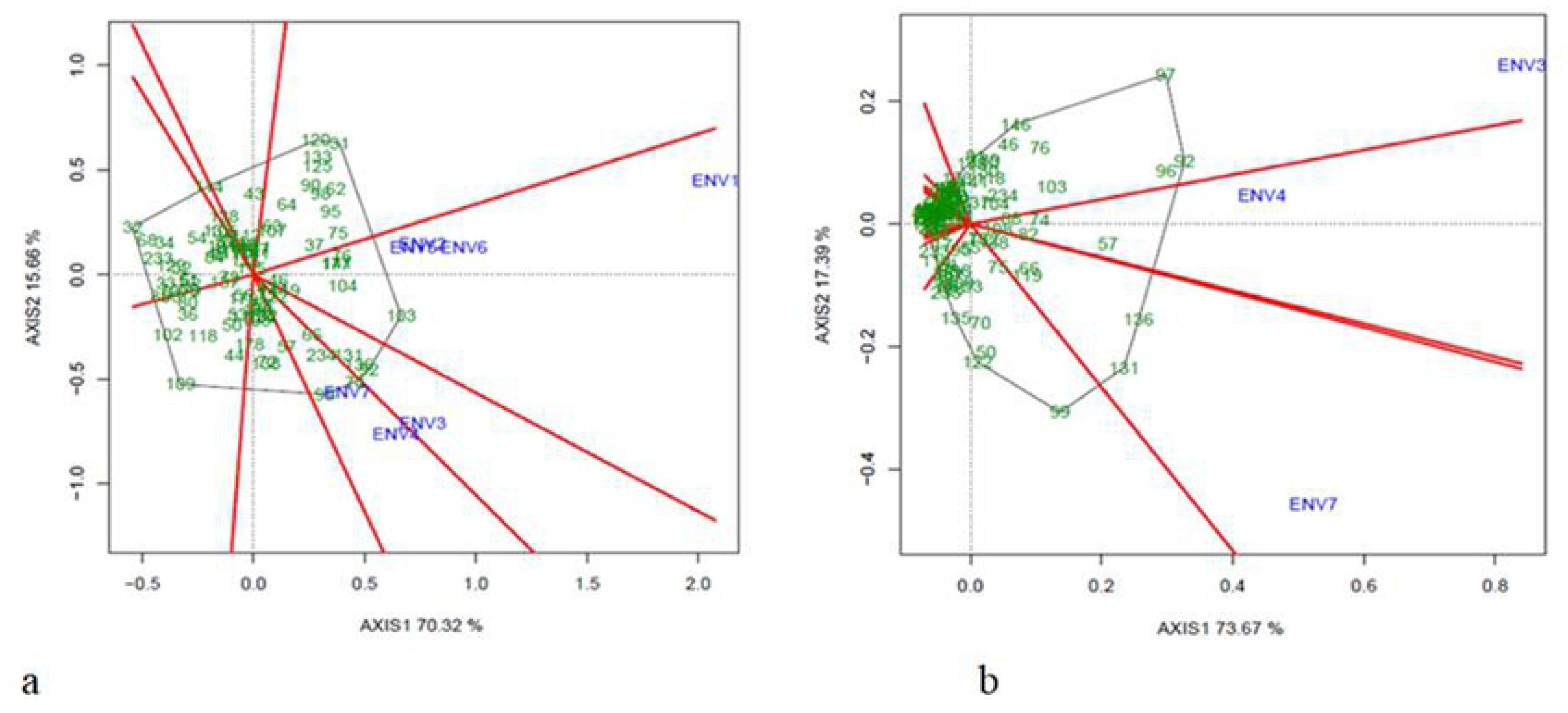
Which Won Where/What graphic of GGE-Biplot analysis for leaves and stem resistance evaluation. (a) GGE-Biplot analysis for phenotype evaluation against *P. infestans* in leaves. (b) GGE-Biplot analysis for phenotype evaluation against *P. infestans* in stem. ENV codes are shown in table 1.

On the other hand, the genotypes CCC003 (32) and CCC101 (103) were identified as extreme resistant and susceptible to late blight, respectively evaluated in leaves, while the genotypes CCC086 (91) and CC133 (131) were identified in stem as extreme resistant and susceptible, respectively. The numbers in the parentheses are the running ones as shown in the Figure 2. According to the rAUDPC, 44 genotypes are resistant in the stem evaluation. For leaves, five genotypes CCC003 (32), CCC051 (68), CCC005 (34), CCC115 (114), and the control Unica (233) were the most resistant.

### 3.4. Stable QTL identified for late blight resistance in leaves and stem

In total 16 organ-specific QTL distributed in nine of the 12 chromosomes were identified (Table 3 and Figure 3). The QTL detected explain from 13.7% to 50.9% of the phenotypic variance (late blight resistance) (Table 3). Six QTL were detected through the leaves evaluation, three under locality Subachoque and the other three for the locality La Unión. Two of these QTL were considered as stable as they were detected in the same organ (leaves) in the two localities, RPiLSub3 and RPiLLU3b for Subachoque and La Union, respectively. The QTL RPiLSub3 explains 25.1% while the QTL RPiLLU3b explains the 24.3% of the resistance to *P. infestans* (Table 3). On the other hand, ten QTL were detected for late blight stem resistance. Five QTL were detected in Subachoque locality and the other five in La Unión. The QTL coded as RPiSLU12b explains the highest phenotypic variance of 50.9% identified through the stem evaluation.

**Figure 3.**
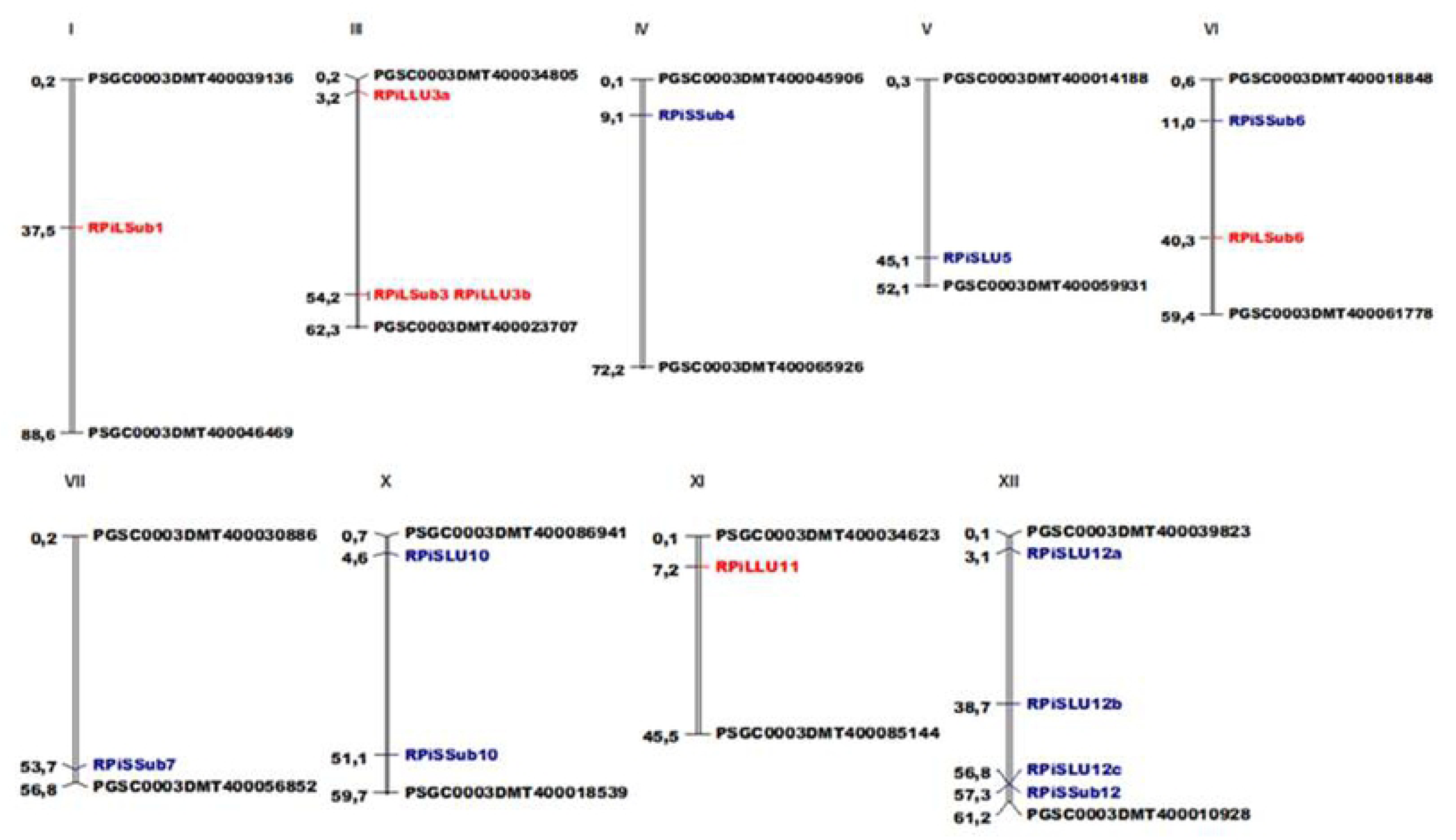
SNPs mapped in the potato chromosomes of Group Phureja. Genomic distances are given in Mbp according to the PGSC V4.03 pseudomolecules [29]. In red are presented the QTL for late blight resistance detected in the phenotypic response for leaves, while in blue are presented those detected for stems. In black are presented the markers that limit each chromosome in the potato genome. Diagram plotted using MapChart software [41].

### 3.5. QTL and candidate genes for organ-specific resistance to late blight

The DNA genome sequences harboring associated SNPs to late blight resistance were analyzed *in silico* using BLAST against the current potato reference genome (v4.04) and to the potato consensus QTL map [29]. Those genes which co-localized with the SNPs associated to late bight resistance were considered as candidate genes for late bight resistance. In total, 15 candidate genes were identified, ten for stem resistance and five for leaves resistance. Among the total of candidate genes, those corresponding to the QTL RPiSSub12, RPiSLU10, RPiSLU12b, and RPiSLU12c have not been previously annotated, while the other 11 candidate genes, have known functional annotation in the current potato genome, with diverse protein domains (Table 3). Interestingly, the QTL RPiSSub4, which explains 17.3% of the resistance to late blight in stem, co-localized with a gene coding for a leucine-rich repeat containing protein kinase, a class of genes shown to be involved in plant immunity.

## 4. Discussion

We identified 16 organ-specific QTL for resistance to *P. infestans* and 15 candidate genes that might have a putative function in the defense response of potato against this pathogen. For that an association panel of 150 accessions from *S. tuberosum* Group Phureja was evaluated under seven environments conditions and genotyped through GBS and 2b-RAD. The locations chosen for the phenotypic evaluation to late blight organ-specific disease response (La Unión-Antioquia and Subachoque-Cundinamarca) are important potato production regions in Colombia with high natural pathogen pressure. Moreover, these are geographic regions where the late blight disease response in the Group Phureja has been evaluated before and the *P. infestans* populations in these regions have been explored [43]. Nevertheless, it will be interesting to explore the actual genetic diversity of the *P. infestans* populations in these two locations, in order to elucidate connections between this variability and the variability found both in the defense response of the evaluated genotypes and also in the identified QTL. Furthermore, this will let directing the potato breeding programs toward the development of resistant adapted materials for these important potato production regions.

In the genetic stability analysis, the addition of the phenotypic data (ENV1, ENV2, ENV5, and ENV6) shows a major variation of the resistance behavior in the genotypes and revealed the presence of Meta Environments. In the phenotypic evaluation of stems, the genotypes behavior was less dissimilar with a clear resistance trend. On the other hand, in the leaves phenotypic evaluation, the MET 1 (Figure 3) grouped the phenotypic data, which were obtained when the climate phenomenon called La Niña affected Colombia, and in consequence those crop cycles presented higher relative humidity than MET 2. While in the MET 2 those environments considered as normal years with an average relative humidity were grouped. Therefore, as previously reported, the environments with higher relative humidity seem to the most favorable for the late blight disease development [5,44].

In this study 16 organ-specific QTL conferring resistance in potato Group Phureja against *P. infestans* were identified, explaining high levels of phenotypic variance or late blight resistance (13.7% to 50.9%) thus these regions could be considered as QTL with major effects [45]. Ten of these QTL were detected through the stem evaluation while the other six QTL were detected through the leaves evaluation. Despite that it is known that the response in a plant to a pathogen can be organ specific [24], according to our understanding this is the first organ-specific late blight resistance QTL detection study through the phenotyping in leaves and stem from *S. tuberosum* Group Phureja.

The organ specific analyses have been considered as a useful way to dissect and understand the plant pathogen polygenic resistance. Detection of organ-specific pathogen resistance QTL has already been published in several crops such as maize (*Zea mays*) [46], tetraploid potato [23–25] and wheat [47]. The resistance responses in different plant organs have been associated with the ability and speed of defense related metabolites production, as well as how these metabolites can be transported to other plant organs and then increase the plant pathogen basal defense. Dannan et al., [23] report that one of the advantages of the high level of resistance to late blight exhibited by the potato stem compared to the leaves, is the ability of the stem to recover the plant leaves and thus could have a significant potential in controlling late blight epidemics in the field.

Several resistance QTL to late blight have been reported in *S. tuberosum* Group Phureja [4,48,49], however, none of those QTL were identified in this study. This could be explained by the high effect of the environments where the phenotypic evaluations were performed. Also, the plant materials employed, their genetic backgrounds, and the phenotyping restricted to specific plant organ could have an important influence in the QTL detection. Nevertheless, we are reporting a stable QTL, which receives its name according to the locality in which it was detected, RPiLSub3 and RPiLLU3b, in Subachoque and La Unión conditions respectively. In Subachoque this QTL explains 25.1% while in La Unión explains the 24.3% of the resistance to *P. infestans*. The phenotypic variance explained by a QTL could change depending on the environment where the phenotypic data were collected and also by the evaluated organ. This variability has been reported in QTL for maize [50], barley [51], and *Brassica rapa* [52].

The genome regions harboring the SNPs associated with resistance to late blight harbor genes coding for proteins carrying diverse domains and were established as candidate genes for resistance to late blight. We propose 15 candidate genes, four of them have not functional annotation so far. Interestingly, two of these anonymous QTL, are those explaining the highest percentage of phenotypic variances, RPiSLU12c and RPiSLU12b, which explain 38.4% and 50.9% of the resistance to late blight, respectively. These genomic regions have a big potential to be explored in the near future for functional validation and establish their annotation.

Both qualitative and quantitative resistance have been described in the *S. tuberosum - P. infestans* interaction [6,18,46,53]. Thus, it is not surprising that within the set of candidate genes was found a gene coding for a leucine-rich repeat protein kinase family, a protein typically related to qualitative resistance, co-localizing with the RPiSSub4 QTL. This finding reinforces the thesis that the qualitative resistance governed by the Avr-R interactions and the quantitative resistance governed by QTL can act simultaneously. On the other hand, the fact that the majority of the identified candidate genes do not belong to classical protein domains of resistance genes, goes accordingly to previous findings where it has been shown that the proteins carrying the classical resistance domains could be just a few of the whole set of proteins that governs the quantitative resistance [22].

Here four candidate genes coding for protein kinases were identified co-localizing with the QTL RPiLLU11, RPiSSub4, RPiSLU5, and RPiSLU12a. The role of kinases has broadly described in plant resistance [54–55]. Their roles have been described as a receptor molecule and also involved in the defense signaling pathway [58]. In *S. tuberosum* some protein kinases have been reported such as calcium-dependent protein kinases (CDPKs) decode calcium (Ca^2+^) signals and activate different signaling pathways involved in hormone signaling and both abiotic and biotic stress responses [59]. In this study, the authors identified different organ-specific expression levels among the genes *CDPKS*. The higher expression of these genes was in roots, stolons, and microtubers compared with the level in leaves and stems (i.e. StCDPK7). Also, the potato CDPKs can be sorted into two classes, those which are ubiquitously expressed in different tissues and those that have specific expression pattern which are approximately one-third of the CDPK family [59].

The results presented here demonstrate the great potential of the Group Phureja as an alternative source of resistance to potato late blight. In fact, the diploid nature of this group facilitates its use in molecular breeding programs, whose aim is the transferring of resistance into tetraploid potatoes gene pool. This article shows that the responses of potato plants to *P. infestans* infection are organ-specific. These differential responses can have an important impact on the behavior of the genotype and therefore on its production. Therefore, these results open the door to continue exploring this type of resistance, so in the near future we can develop strategies that integrate the specific responses in the potato improvement and production programs.

It is known that the dynamics with which effectors and *R* genes interact in resistant potato genotypes versus susceptible genotypes differ in the level of transcription, as well as in the speed with which it is induced in tuber [25] and in leaves [60]. Now, our results open the way for studies of expression demonstrating the above in potato leaves of the Group Phureja and for the first time in potato stem.

The identified QTL in this research are an advance in the discovery of new organ-specific genetic factors involved in the resistance response of *S. tuberosum* Group Phureja to *P. infestans* infection. Also, the genomic regions associated and the candidate genes proposed are resistance sources to be functionally validated in the near future. The environment specific QTL here described hides interesting molecular mechanisms to be explored to a better understanding the genotype-environment interactions in the potato - *P. infestans*interaction. Finally, the results could be integrated into potato genetic breeding programs focused on marker assisted selection, as well as, for the development of local adapted new cultivars or by combining or pyramiding these QTL to develop environment independent resistant cultivars.

## 6. Author’s contributions

DJ: Designed and established the experiments, took the data, analyzed the results, prepared the draft of manuscript. JS: Revised and analyzed the results, prepared the draft of manuscript. AB: Contributed with conceptualization of the original idea, revised the results, analyzed the results, revised the draft of manuscript. JL: Revised and analyzed the results, revised the draft of manuscript. TM: Conceptualized the original idea, supervised the whole research, designed the experiments, revised the results and analyses and revised the manuscript.

## 8. Acknowledgments

We are grateful to Dr. Klaus Dehmer for supporting this work with plant material consisting in diploid accessions from the *IPK Germplasm Bank of Germany* (Leibniz-Institut für Pflanzengenetik und Kulturpflanzenforschung. We would like to thank to Jorge Duitama from “Universidad de los Andes” for the bioinformatics support; to Diana Duarte, Boby Mathew, and Said Dadshani from INRES Institute of University of Bonn for their scientific support and to María Cecilia Delgado for her collaboration in the establishment of the experiments in field conditions.

## 9. Supporting information captions

**SI 1.** a. Analysis of variance for genotype, environment and genotype x environment among the association panel for a. leaves phenotyping. b. stem phenotyping (p<0.001).

**SI 2.** Transitions and transversion SNPs from the genotyping matrix of 83,862 SNPs.

**SI 3.** rAUDPC values by genotype for leaves and stem resistance evaluations.

**SI 4.** GBS and 2b-RAD genotyping data.

**Conflict of Interest Statement:** The authors declare that the research was conducted in the absence of any commercial or financial relationships that could be construed as a potential conflict of interest.

## 10. Data Availability Statement

The genotyping data supporting the conclusions of this manuscript is available in SI 4.

